# Local population genetic structure of *Andrena flavipes* (Hymenoptera, Andrenidae), a nest-philopatric communal mining bee, reveals isolation-by-distance and high inbreeding

**DOI:** 10.64898/2026.10.14.759052

**Authors:** Lotte F. Wösten, Frank F.M. Becker, Joost van den Heuvel, Bart A. Pannebakker

## Abstract

Wild bee populations are declining globally and understanding how species-intrinsic traits like nest-philopatry shape local population genetic structure is increasingly important for their conservation. However, few studies have explored this at local spatial scales. To fill this gap, we used SNPs in the nuclear vitellogenin (Vg) gene to assess the local population structure and genetic diversity of the philopatric yellow-legged mining bee, *Andrena flavipes*, in the Netherlands, one of the most fragmented and urbanized countries in Europe. We investigated whether its reported nest-philopatry, combined with a fragmented landscape, would lead to genetic differentiation and isolation-by-distance (IBD) between populations, and low genetic diversity and inbreeding within populations. A total of 144 *A. flavipes* bees were sequenced for the full-length Vg gene and genotyped for SNPs. Despite high overall genetic variation, we found a heterozygote deficit and positive inbreeding coefficient in most populations, suggesting non-random mating. Low genetic differentiation was detected between populations, with no distinct genetic clusters, but with a pattern of IBD. These results suggest extensive but spatially structured gene flow, consistent with philopatric behaviour, and indicate that increasing isolation between nest-aggregations potentially leads to genetic differentiation over time. Our findings demonstrate that even in a common and widespread species, nest-philopatry and landscape fragmentation can drive inbreeding and spatially structured gene flow at local scales, highlighting the importance of maintaining habitat connectivity and semi-natural elements in agricultural and urbanized landscapes to sustain wild bee populations.

## Introduction

Pollinators ensure the entire reproduction and survival of 88 % of wild flowering plants and around 35 % of global food production (Klein et al. 2007; Ollerton et al. 2011), making them a critical group in terrestrial ecosystems (Cardoso et al. 2020). Among the pollinators, bees are thought to be the major pollinators of wild plants and agricultural crops (Potts et al. 2010). In particular, the western honeybee (*Apis mellifera)* is generally regarded as the most important bee species in terms of pollination and has also been the most extensively studied bee (Potts et al. 2010). However, during the last decade, research has pointed out that pollination services of wild bees have been underestimated (Klein et al. 2007; Breeze et al. 2011; Garibaldi et al. 2011) and that wild bees might even be more efficient pollinators than honeybees (Holzschuh et al. 2012; Garibaldi et al. 2013; MacInnis & Forrest 2019). Moreover, studies have shown that (wild) bee species diversity increases pollination efficiency and effectiveness resulting in increased fruit and seed production in plants (Greenleaf & Kremen 2006; Albrecht et al. 2012; Brittain et al. 2013; Blitzer et al. 2016).

However, there is evidence of wild bee decline globally (Zattara & Aizen 2021). This decline is largely driven by habitat loss and fragmentation, but pressures like climate change, agrochemicals and invasive species also contribute to this decline (Potts et al. 2010; LeBuhn & Vargas Luna 2021). Many of these pressures result from intense human land-use practices, like urbanization and agriculture, which alter the habitat suitability for wild bees and consequently their survival, persistence and movement across landscapes (Klein et al. 2007; Winfree et al. 2009; Ollerton et al. 2014). Habitat loss can lead to smaller population sizes and a loss of genetic diversity in wild bees (Packer & Owen 2001; Zayed et al. 2005), while habitat fragmentation increases the distance between suitable habitat patches, restricting dispersal and gene flow, which can amplify the negative genetic effects of inbreeding and genetic drift, leading to higher genetic differentiation between wild bee populations (Fischer & Lindenmayer 2007; Jha & Kremen 2013b; Jha 2015). Wild bees may be especially vulnerable to habitat loss and fragmentation due to their haplodiploid genetic system, which can reduce their effective population size (*N_e_*) compared to diploid species of equivalent census size (Hedrick & Parker 1997). Furthermore, many bees have complementary sex determination (CSD) (Cook 1993). Heterozygosity at the CSD locus results in females, while hemizygosity or homozygosity results in males (Whiting 1943; Cook 1993). Consequently, in small, inbreeding bee populations, sterile diploid males can be formed, which can lead to decreased population growth, an even further reduced *N_e_* and inbreeding depression, initiating a rapid extinction vortex known as the diploid male vortex (Zayed 2004; Zayed & Packer 2005; Leung & van der Meulen 2022).

The consequences of habitat loss and fragmentation for wild bees can be further aggravated by nest-philopatry, the tendency of individuals to stay in or return to their birthplace to reproduce. Many wild bee species are philopatric and nest in aggregations, which can result in nest-aggregations persisting for many generations (Danforth et al. 2019). When bees exhibit strong nest-philopatry, these aggregations may become increasingly isolated, which can lead to high genetic differentiation between populations and to a pattern of isolation-by-distance (IBD) (Stow et al. 2007; Zayed & Packer 2007; Davis et al. 2010). Such isolation makes populations more susceptible to loss of genetic diversity through inbreeding and genetic drift, increasing the risk of inbreeding depression which can ultimately lead to extinction (Wright 1948; Gilpin & Soulé 1986; Bohonak 1999). While some cavity-nesting bees can profit from increased urbanization due to increased suitable nesting sites in human-made structures (Cane et al. 2006; Vickruck & Richards 2017), ground-nesting bees, comprising the majority of wild bees (75 %) (Michener 2007; Harmon-Threatt 2020; Antoine & Forrest 2021) are mostly not positively affected (Matteson et al. 2008; Fortel et al. 2014; López-Uribe et al. 2015).This is because in urban and agricultural landscapes, the soil is heavily modified and the proportion of bare soil dramatically reduced, limiting suitable nesting sites (Jha & Kremen 2013a). As the nesting substrate chosen by females is species specific (Cane 1991; Antoine & Forrest 2021), successful development of ground nesting-bees occurs only when females nest in soils that meet the species’ ecological requirements (Harmon-Threatt 2020). Therefore, when suitable nesting sites are limited or unevenly distributed, the availability of these resources may also constrain population size and geographic range of wild bees (Vickruck & Richards 2017). In certain ground-nesting bee species, nest-philopatry and within-nest mating have been found to result in high levels of inbreeding (Paxton et al. 1996; Danforth et al. 2003), which could be further amplified in fragmented landscapes.

The signs of dispersal, inbreeding, and genetic drift can be detected in a population’s genetic structure, which reflects the species’ demographic history and gene flow between populations over time and space (Slatkin 1985; Bohonak 1999), providing insight into species’ dispersal and colonization patterns (Bohonak 1999; Broquet & Petit 2009). Population structure can help to reveal environmental factors that either facilitate or hinder movement across landscapes at various spatial scales. Many population structure studies have historically focused on large spatial scales, such as global, continental, or regional scales, particularly for vertebrates and other large-bodied species, where gene flow patterns are easier to observe across these larger scales (Ballare & Jha 2021). However, for small-bodied species that have limited dispersal abilities either by species intrinsic traits (like philopatry or limited flight capacity) and/or environmental constraints (like habitat fragmentation or soil-specificity), it is more useful to study population structure at finer, local scales. When dispersal is limited, allele frequencies tend to change over short distances, so when sample units are too large relative to dispersal distance, subpopulations might be merged that are in fact genetically distinct, thereby losing resolution and obscuring fine-scale genetic differentiation (Bluher et al. 2020). Examining population structure at a local scale, might yield more insight into the current dispersal and nesting dynamics of wild bee species and their susceptibility to environmental change (Bluher et al. 2020; Ballare & Jha 2021). This population-level information can then be used to design feasible and effective conservation strategies to protect and support wild bee populations, which are urgently needed (Dorian et al. 2024).

An interesting wild bee species to study the effects of nest-philopatry on population genetic structure and genetic diversity is *Andrena flavipes* (Hymenoptera, Andrenidae), a solitary ground-nesting bee that is widely distributed throughout the Palearctic region and commonly found in urban and rural landscapes. It is polylectic on 13 plant families (Alfken 1913; Perkins 1919; Chambers 1968; Westrich 1989a, b) and one of the most important crop pollinators in Europe (Kleijn et al. 2015). It has two overlapping generations per year (Alfken 1913; Westrich 1989b; Peeters et al. 2012) and is easily recognizable in the field, making it an ideal study species. Despite being solitary, *A. flavipes* females often nest in aggregations that can contain up to thousands of individuals (Westrich 1989a, b; Hallmen 1993; Peeters et al. 2012), and can persist for many generations (Chambers 1968; Boyes et al. 2024). These aggregations are not organized colonies as those in eusocial bees, but rather a form of communal nesting. Females prefer to dig their nest in firm, clayey or loamy soil with sparse vegetation, preferably on a sun-exposed slope (Alfken 1913; Westrich 1989a, b; Hallmen 1993; Peeters et al. 2012). Males prefer to mate with females from within their own natal nest-aggregation (Butler 1965; Peeters et al. 2012) and females tend to only mate within, and to nest within, their natal nest-aggregation (Butler 1965). Moreover, *A. flavipes* nest-aggregations have been found to be restricted in size and colonization of new nest-aggregation seems to occur rarely (Butler 1965). Taken together, these traits may result in genetic isolation, high differentiation and inbreeding among closely situated nest-sites of *A. flavipes*.

Previous studies on the population genetic structure of *A. flavipes* have been done on large spatial scales only, for instance in Austria using microsatellites (<400 km) (Rupprecht 2022), and in Portugal (<250 km) (Henriques 2021), the Iberian Peninsula (<800 km) (de Sousa 2021; de Sousa et al. 2023) and Europe (<2500 km) (Ortego et al. 2024) using SNPs. However, since bees are in general very small and are not expected to disperse over long distances (Ballare & Jha 2021) it is crucial to investigate the local population structure (∼10 km), to gain insight into the dispersal patterns and nesting dynamics of *A. flavipes.* Furthermore, of the few studies that looked at population structure in other wild bee species at local scales, strong philopatric tendencies have been shown to lead to population structure at very small scales, even though substantial gene flow was detected at larger scales (López-Uribe et al. 2015; Schenau & Jha 2017; Ballare & Jha 2021).

In this study, SNPs in the nuclear vitellogenin (Vg) gene were used to assess the local population structure and genetic diversity within and between five populations of *A. flavipes* in an area of <13 km in the Netherlands, one of the most fragmented and urbanized countries in Europe (European Environment Agency et al. 2011; European Environment Agency & Swiss Federal Office for the Environment 2016). Given the evidence of high genetic variation and population-level signal in the Vg gene in honey bees (Ilyasov et al. 2022; Leipart et al. 2022), we explored its utility in *A. flavipes*. We hypothesized that based on *A. flavipes’* reported 1) nest-philopatry, 2) restriction in nest-site size and 3) rare nest-colonization events, combined with a fragmented landscape, this will lead to genetic differentiation between populations, low genetic diversity within populations and a pattern of IBD.

## Material and Methods

### Specimen collection

Between April-May of 2022 and April-July 2023, *A. flavipes* bees were net-collected from five locations consisting of 18 nest-sites in the Southern Gelderse-Vallei (provinces of Gelderland and Utrecht in the Netherlands) (Fig. 1; Fig. S1; Table S1). Additionally, three female *A. flavipes* bees were net-collected from three locations outside of the Gelderse Vallei area, in Gemert and Handel (Province of Noord-Brabant in the Netherlands) and Maastricht (Province of Zuid-Limburg in the Netherlands) (Table S1). The three bees collected from Gemert, Handel and Maastricht were used as outgroup in this study. Field collection trips were made based on the database of waarneming.nl (https://waarneming.nl/), where sightings of *A. flavipes* have been recorded since 1999. Sightings after 2010 and with five or more individuals at one nest-site were prioritized.

**Figure 1.**
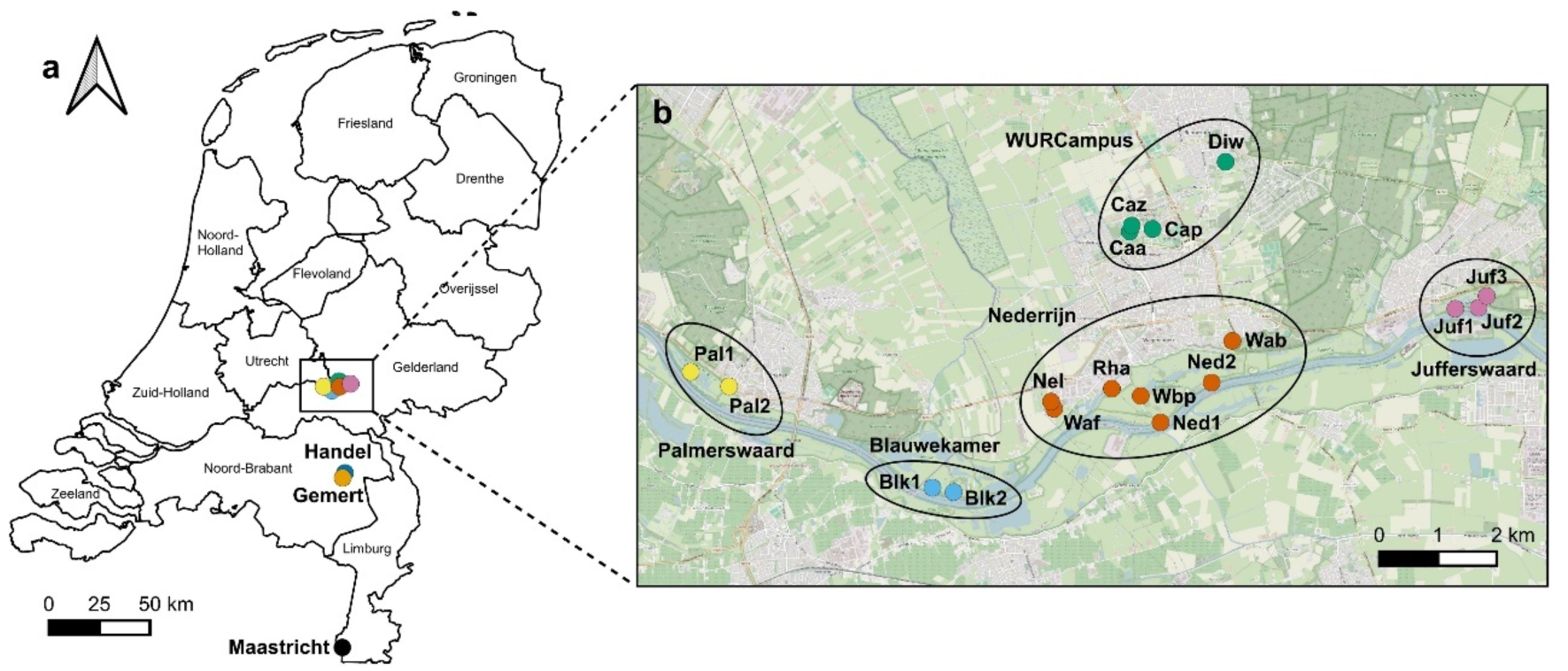
Map of nest-sites and populations of net-collected *A. flavipes* bees in the Netherlands. **(a) Map of five populations from the Southern Gelderse-Vallei and three outgroup populations in the Netherlands.** Names of the eight populations are Palmerswaard; BlauweKamer; Nederrijn; WURCampus; Jufferswaard; Handel; Gemert and Maastricht. The map indicates the names of the seven Provinces of the Netherlands. **(b) Map of nest-sites that make-up the five populations from the Southern Gelderse-Vallei.** Names of the 21 nest-sites are Palmerswaard 1(Pal1); Palmerswaard 2 (Pal2); Blauwe Kamer 1 (Blk1); Blauwe Kamer 2 (Blk2); Nel Vietsstraat (Nel); Wageningse Afweg (Waf); Rijnhaven (Rha), Wageningse Bovenpolder (Wbp); Nederrijn 1 (Ned1); Nederrijn 2 (Ned2); Wageningse Berg (Wab); Campus Aurora (Caa); Campus Zodiac (Caz); Campus Plaza (Cap); Diedenweg (Diw); Jufferswaard 1 (Juf1); Jufferswaard 2 (Juf2), Jufferswaard 3 (Juf3), Handel, Gemert and Maastricht. The map was made using *QGIS v3.38* (Free Software Foundation, Inc., Boston, MA, USA).

All the net-collected bees were stored in 1.5 mL microfuge tubes, frozen in liquid nitrogen and stored at -80°C for subsequent analyses. Species identification of the bees was done morphologically and genetically, using a stereoscope and COI DNA barcoding (see section COI amplification and species identification).

### DNA extraction

Genomic DNA (gDNA) was extracted from the head and thorax of each bee, using the DNAeasy^®^ Blood and Tissue kit (Qiagen, Venlo, The Netherlands), following the manufacturer’s protocol (Fig. S1). The head and thorax were placed in a 1.5 mL microcentrifuge tube and grinded in liquid nitrogen using a pestle. After grinding the tissue, 180 μL ATL buffer and 20 μL proteinase K were added and the samples were incubated overnight at 56°C in a thermo-shaker at 350 rpm. After extraction, the gDNA was eluted with 100 mL AE buffer, and stored at -20°C.

### COI amplification and species identification

Species identification was done on all samples using DNA barcoding with the mitochondrial marker cytochrome *c*-oxidase subunit I (COI) (Fig. S1). The primer pair BeeCox1F1 (5’-TAGTCAACAAATCATAAAGATATTGG-3’) and BeeCox1R2 (5’-CCAAATCCTGGTAGAATTAAAATATA-3’), with an expected PCR amplicon size of 670 bp, was used to amplify and sequence the COI gene of *A. flavipes* (Bleidorn & Henze 2021). The PCR was performed in standard 0.2 mL PCR tubes, wherein each PCR reaction contained 1 μL of extracted gDNA (<250 ng), 5 μL 5X GoTaq^®^ G2 Green buffer (Promega, Madison, Wisconsin, USA), 1 µL (5 mM) dNTP’s, 1 μL (10 μM) forward primer, 1 μL (10 μM) reverse primer, 0.125µL GoTaq^®^ G2 DNA Polymerase (Promega, Madison, WI, USA) and 15.875 µL Ambion™ Nuclease-Free Water (Invitrogen, ThermoFisher Scientific, Waltham, MA, USA) to a final volume of 25 µL. A negative control was included for each PCR that was run. The PCR cycle was as follows: initial denaturation at 94°C for 2 min, 40 cycles of 94°C for 30 sec, 50°C for 45 sec, 72°C for 1 min, and then final extension at 72°C for 10 min and hold at 12°C. 4 µL of each PCR amplicon was checked on a 1% TAE agarose gel to verify amplification and absence of contamination. Subsequently, the PCR amplicons were sent off for Sanger sequencing (Eurofins Genomics, Ebersberg, Germany) and sequenced in the forward direction. The quality of the COI sequences was examined using Geneious Prime v2023.2.1 (Dotmatics, Boston, MA, USA). NCBI-BLAST (Sayers et al. 2022) was used to identify each of the obtained COI sequences. A median-joining haplotype network was constructed from the COI sequences using *PopART v1.7* (Leigh & Bryant 2015) to visualise the relationship between COI haplotypes across populations.

### *de novo* genome sequencing and assembly

Following correct species identification, a *de novo* genome assembly was made using single male *A. flavipes* net-collected from the Nederrijn, Wageningen, Gelderland, the Netherlands (Fig. 2; ID: Ned2, 51°57’35.4"N 5°40’46.4"E). DNA was extracted using the high-molecular-weight DNA extraction protocol for long-read sequencing (Russo et al. 2022).

**Figure 2.**
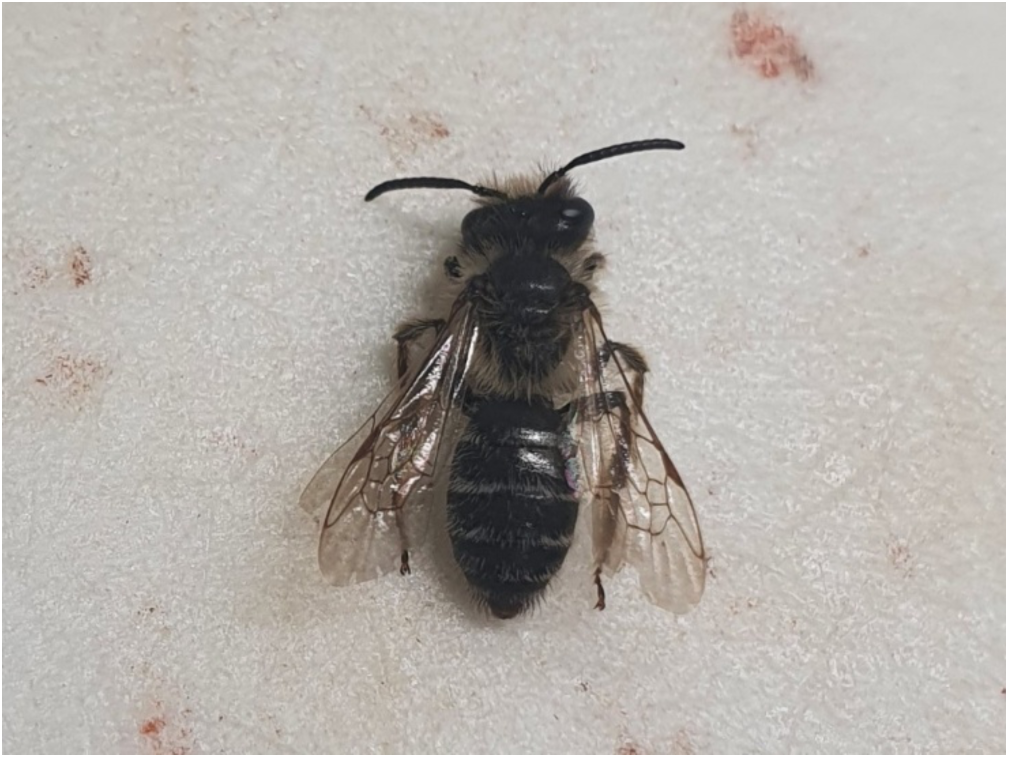
Image of the *A. flavipes* male (Aflav121_Ned2) used for *de novo* genome sequencing. Coordinates nest-site (Ned2): 51°57’35.4"N 5°40’46.4"E.

Subsequently, library preparation was done using the Ligation Sequencing kit (SQK-LSK114, Oxford Nanopore Technologies, Oxford, UK) following the manufacturer’s protocol. The final library was quantified on the DeNovix DS-11 FX (DeNovix, Wilmington, DE, USA) using the DeNovix dsDNA High Sensitivity Assay kit following the manufacturer’s protocol (TN144, DeNovix, Wilmington, DE, USA). After adapter ligation, the flow cell (FLO-MIN114 (R10.4.1), Oxford Nanopore Technologies, Oxford, UK) was primed using the Flow Cell Priming kit (EXP-FLP0002, Oxford Nanopore Technologies, Oxford, UK) following the manufacturer’s protocol, after which the final prepared library was loaded.

Sequencing was performed using the MinION™ Mk1C device (Oxford Nanopore Technologies, Oxford, UK) and the MinKNOW v23.07.12 software was used to operate sequencing. Post-run high accuracy base calling (HAC) was performed on the raw pod5 files downloaded from the MinION server using the data processing toolkit Dorado v0.4.2 (Oxford Nanopore Technologies, Oxford, UK), which resulted in the conversion of pod5 files to fastq files, with a minimum qscore of 10. Read quality statistics were generated using NanoPlot (De Coster & Rademakers 2023).

The reads were assembled using *Flye 2.9.1* (Kolmogorov et al. 2019). The assembled genome was assessed for completeness with Benchmarking Universal Single-copy Orthologs (BUSCO 6.0.0) using the arthropoda_odb10 database (Manni et al. 2021; Tegenfeldt et al. 2024).

### Vitellogenin amplicon sequencing

#### Vitellogenin primers and amplification

PCR primers were developed using Primer3 in Geneious Prime v.2023.2.1 to amplify the whole Vg gene (including introns) of *A. flavipes* (Table 1; Fig. S1). The full-length Vg gene of the honeybee (*Apis mellifera*), spanning positions 5,029,433 to 5,035,683 on NC_037641.1 of the genome assembly Amel_HAv3.1 (Wallberg et al. 2019), was used to find the orthologous Vg gene in the *de novo* assembled genome of *A. flavipes* (Laboratory of Genetics, WUR, Netherlands). In addition, the Vg primers were extended with tails (provided in EXP-PBC096 protocol, Oxford Nanopore Technologies, Oxford, UK) incorporated into the 5’ ends of the Vg primers, so that barcodes unique for each individual bee could be attached to the Vg sequences.

**Table 1.**
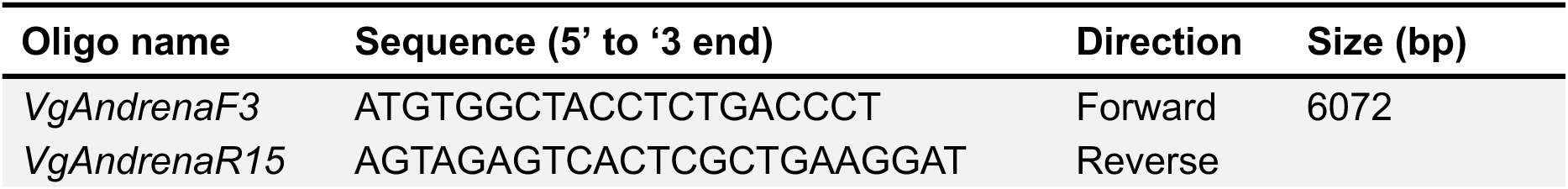
Primers used for the amplification of the putative nuclear Vg gene of *A. flavipes*.

A tailed PCR was performed in two 96-well plates, wherein each PCR reaction contained 1 µL of extracted gDNA (<100 ng), 12.5 µL 2X concentrated PCRBIO VeriFy™ Mix (PCRBIO VeriFi™ Polymerase, 6 mM MgCl2, 2 mM dNTPs, enhancers and stabilizers) (PCR Biosystems, London, UK), 1 µL (10 µM) forward primer, 1 µL (10 µM) reverse primer and 9.5 µL of Ambion™ Nuclease-Free Water to a final volume of 25 µL. The cycling conditions were as follows: initial denaturation at 95°C for 1 min, 34 cycles of 95°C for 15 sec, 68°C for 15 sec, 72°C for 3 min, and then final extension 72°C for 5 min and hold at 12°C. A previously verified sample (Aflav_120) and a negative control (Nuclease-Free Water) were included in the two PCR plates that were run. 1 µL of each PCR reaction, including the positive and negative control, was checked on a 1% TAE agarose gel to verify amplification and the lack of contamination. Each PCR reaction (24 µL) was cleaned using 24 µL SPRI beads (SpeadBead Magnetic Carboxylate Modified Particles, Cytiva, Marlborough, MA, USA) and washed two times with 100 µL 70 % EtOH before being eluted in 20 µL Nuclease-Free Water. After clean-up of the PCR plates, the concentration of each amplicon was measured in the SPECTRAFluor Plus Microplate Reader (Tecan, Männedorf, Zwitserland) with XFluor v4.5.1 using the Quant-iT™ PicoGreen^®^ dsDNA Assay kit (Invitrogen ThermoFisher Scientific, Waltham, MA, USA) following the manufacturer’s protocol. For each PCR reaction, the DNA concentration was calculated, with values ranging from 0 ng/µL to 45 ng/µL in plate 1 and 1 ng/µL to 82 ng/µL in plate 2. Samples with DNA concentrations less than 2 ng/µL were discarded.

#### Library preparation

The barcodes (BC01-BC96) provided in the PCR Barcoding Expansion 1-96 kit (EXP-PBC096, Oxford Nanopore Technologies, Oxford, UK) were used to barcode the *A. flavipes* samples (Fig. S1). Barcoding PCR was performed in two 96-well plates, wherein each PCR reaction contained 0.3 µL (10 µM) barcode, 7.2 µL (0.5 nM) cleaned PCR product and 7.5 µL LongAmp^®^ Taq 2X Master Mix (New England BioLabs, Ipswich, MA, USA) to a final volume of 15 µL. The cycling conditions for the barcoding PCR were as follows: initial denaturation 95°C for 3 min, 15 cycles of 95°C for 15 sec, 62°C for 15 sec, 65°C for 6 min, and then final extension at 65°C for 6 min and hold at 4°C. After the barcoding PCR, the concentration of each amplicon was measured again in the SPECTRAFluor Plus Microplate Reader with XFluor v4.5.1 using the Quant-iT™ PicoGreen^®^ dsDNA Assay kit following the manufacturer’s protocol. Samples with DNA concentrations less than 24 ng/µL for plate 1 and 26 ng/µL for plate 2 were discarded. A total of 30 µg of barcoded samples within each of the plates was pooled in a 1.5 mL Eppendorf DNA LoBind^®^ tube (equalizing the concentrations, with the highest concentration set to 1 µL within each plate) (Eppendorf, Hamburg, Germany), creating two barcoded pools of DNA. These pools were cleaned with SPRI beads, adding the same amount of SPRI beads as the volume of the pooled DNA and washed two times with 200 µL 80 % EtOH, before being eluted in 200 µL Ambion™ Nuclease-Free Water. The DNA concentration of the two cleaned barcoded DNA pools was measured on the DeNovix DS-11 FX using the DeNovix dsDNA High Sensitivity Assay kit following the manufacturer’s protocol (TN144). Afterwards, 1.5 µg of each barcoded and cleaned DNA pool was prepared in 48 µL nuclease free water. Subsequently, library preparation was done using the Ligation Sequencing kit (SQK-LSK114, Oxford Nanopore Technologies, Oxford, UK) following the manufacturer’s protocol (Fig. S1).

#### Nanopore sequencing

Sequencing and base calling were performed as described above for de novo sequencing, with the following differences: we washed the flow cell with the Flow Cell Wash kit (EXP-WSH004, Oxford Nanopore Technologies, Oxford, UK) following the manufacturer’s protocol between the two sequencing runs, and after the base calling step we demultiplexed the reads into separate files for each barcode.

#### Bioinformatic analysis

To create Vg haplotype consensus sequences for each demultiplexed sample, two different methods were used for males and females (Fig. S2). Because males are haploid (single Vg haplotype), their reads were assembled directly de novo, whereas the diploid females required variant calling and haplotype phasing to recover both Vg haplotypes.

For creating the haplotype consensus sequences in males, the fastq files were first filtered by size with *seqtk* v1.4 (Li 2023), only selecting fastq files with a length between 5500 and 7000 bp. After filtering for length, the fastq files were randomly shuffled and 40 random fastq files were selected to be assembled with *Flye v.2.9.3* (Kolmogorov et al. 2019) with genomic size set to 6000 and number of iterations set to 10. Subsequently, the resulting Vg sequence assemblies in fasta format were aligned using *Muscle* (Edgar 2004) in *MEGA11* (Tamura et al. 2021) and trimmed in order to get the full-length Vg sequence for each male.

For creating the haplotype consensus sequences in females, the fastq files were filtered by size in python using *biopython* (Cock et al. 2009), only selecting fastq files with a length between 5500 and 7000 bp. The filtered reads were mapped to the de novo genome of *A. flavipes* using *minimap2* (Li 2018) and sorted and indexed with *SAMtools* (Danecek et al. 2021) to create bam and their corresponding bai files for each female sample. In addition, *SAMtools* was used to filter out incomplete and secondary alignments for each female sample. Variant calling was done using *Clair3* (Zheng et al. 2022) using the pileup-method. Afterwards, *Whatshap v.2.3* (Martin et al. 2016) was used to phase the two haplotypes for each of the females. To get the two Vg haplotype sequences for each female sample in fasta format, the generated phased vcf files were filtered to get two haplotype vcf files per female. Subsequently, *GATK v4.3.0* (McKenna et al. 2010) was used to create two alternative reference sequences per female by combining the reference genome of *A. flavipes* with the two haplotype vcf files. Lastly, *SAMtools* was used to filter out the two Vg sequences for each female in the generated alternative reference sequences. The Vg sequences of the females were combined with the Vg sequences of the males resulting in a total of 177 Vg sequences. The 177 Vg sequences were aligned in *MEGA11* (Tamura et al. 2021) using *Muscle* (Edgar 2004), and SNPs were manually curated.

### Vitellogenin annotation

*DnaSP v6.12.03* (Rozas et al. 2017) was used to identify the amount of unique Vg sequences (introns and exons) from the 177 aligned Vg sequences. Afterwards, the online tool *AUGUSTUS* (https://bioinf.uni-greifswald.de/augustus/submission.php, accessed 08-05-2024) was run using *A. mellifera* as the reference organism, reporting genes on both strands with few alternative transcripts. Next, the generated coding sequences were translated in *MEGA11* (Tamura et al. 2021) and the amino acid changes observed in each Vg sequence were manually curated to identify the number of non-synonymous SNPs (nsSNPs) and synonymous SNPs (sSNPs).

### Vitellogenin datasets

Three different datasets were generated for the Vg gene. The first dataset consisted of 177 aligned Vg sequences of 5874 bp in which females were considered as two haploid individuals, so that each Vg haplotype sequence was considered as one individual. The second dataset consisted of SNP genotype data of only females (*n*=33), where each female was considered as a single diploid individual, having two different alleles. The third dataset consisted of SNP genotype data of males and females, where each female was considered as two haploid individuals. Dataset 1 was used for sequence-based analyses, dataset 2 for diploid heterozygosity and deviations from Hardy-Weinberg equilibrium (HWE), and dataset 3 for Principal Component Analysis (PCA) (see below).

### Genetic diversity

Several different genetic parameters were estimated for the Vg gene (*H_o_*, *H_e_*, *F_IS_*, π, *S*, *K*, θ_W_, Tajima’s D) to characterize the genetic diversity of *A. flavipes*.

The expected heterozygosity (*H_e_*) was calculated per population using the *poppr* function in the R package *poppr v2.9.6* (Kamvar et al. 2014). The observed heterozygosity (*H_o_*) and inbreeding coefficient (*F*_IS_) were calculated per population using the *basic.stats* function in the R package *hierfstat v0.5-11* (Goudet 2005). For populations with only one female *H_e_*, *H_o_* and *F_I_*_S_ could not be calculated. The 95 % confidence intervals for *F*_IS_ per population were determined using the *boot.ppfis* function in *hierfstat* with 999 bootstrap replicates to assess significance of *F_IS_* estimates at 0.05 significance level. Deviations from Hardy-Weinberg Equilibrium (HWE) per SNP per population were calculated using the *hw.test* function in the R package *pegas v1.3* (Paradis 2010), with significance assessed using a Monte Carlo test with 999 permutations and a Bonferroni correction to account for multiple testing.

Nucleotide diversity (π), number of segregating sites (S), number of nucleotide differences (K), Watterson’s theta (θ_W_), and Tajima’s D (from total number of mutations) per population and deviations from linkage equilibrium between each SNP pair were calculated using *DnaSP v6.12.03* (Rozas et al. 2017). For the outgroup populations, π, S, K, θ_W_, and Tajima’s D were not calculated. The 95 % confidence intervals for Tajima’s D per population were determined using coalescent simulations with 999 replicates in *DnaSP* under the Standard Neutral Model (SNM) with recombination set to intermediate level to assess significance of Tajima’s D estimates at 0.05 significance level. The significance of deviations from linkage equilibrium for each SNP pair were tested using a Chi-squared test with a Bonferroni correction to account for multiple testing in *DnaSP*. The linkage decay was visualized using the R package *ggplot2 v3.5.1* (Wickham 2016).

### Population genetic structure

To measure genetic differentiation between the populations, the pairwise fixation index (*F_ST_*) was calculated using *Arlequin v3.5.2.2* (Excoffier & Lischer 2010). Before performing population genetic analyses in *Arlequin*, the aligned Vg sequence dataset was first converted to arp format (required file format for *Arlequin*) using DNAcollapser in the online tool *FaBox v1.61* (https://users-birc.au.dk/palle/php/fabox/dnacollapser.php, accessed on 16-05-2024). Pairwise *F_ST_* was calculated with conventional F-statistics (haplotype frequencies only) set as distance, which follows the Weir and Cockerham estimation (Weir & Cockerham 1984). The significance for the pairwise *F_ST_* results was tested at the 0.05 significance level using 999 permutations.

After calculating pairwise F_ST_, a Mantel test was performed using *Arlequin v3.5.2.2* (Excoffier & Lischer 2010), to examine whether there was population structure as a result of geographic distance (Isolation-by-distance; IBD), The pairwise *F_ST_* values were transformed into Slatkin’s Linearized *F_ST_* ((*F_ST_* / (1-*F_ST_*)) (Slatkin 1995; Rousset 1997), which served as genetic distance. Pairwise Euclidean distances between populations were used as geographic distances. As the populations consisted of multiple sites, the weighted mean coordinate of each population was calculated using the *weighted.mean* function in base R, with each site’s coordinate weighted by the number of individuals collected at that site (Table S1). The Mantel test was performed using 999 permutations. Subsequently, Slatkin’s linearized *F_ST_* was plotted against geographic (Euclidean) distance (km) using the R package *ggplot2 v3.5.1* (Wickham 2016).

An Analysis of Molecular Variance (AMOVA) was conducted for the Vg gene using *Arlequin v3.5.2.2* (Excoffier & Lischer 2010) to see how the genetic variance was distributed between and within the populations. The AMOVA was performed using the standard AMOVA setting in haplotypic format with conventional F-statistics (haplotype frequencies only) and the populations considered as one group. The significance for the AMOVA results was tested at the 0.05 significance level using 999 permutations.

A Principal Component Analysis (PCA) was performed for the Vg gene to visualise genetic clustering using the *dudi.pca* function in the R package *ade4 v1.7-22* (Thioulouse et al. 2018). The principal component coordinates (PC1 and PC2) for each individual haplotype were grouped by population. For each group, the centroid (mean position), standard deviation, and sample size was calculated with the R package *dplyr v1.1.4* (Wickham et al. 2023). Standard errors were then computed by dividing the standard deviations by the square root of the sample size. The PCA with centroids and standard errors was visualized using the R package *ggplot2 v3.5.1* (Wickham 2016).

## Results

A total of 114 male and 35 female (N=149) *A. flavipes* were net-collected from five populations consisting of 18 nest-sites in the Southern Gelderse-Vallei (Province of Gelderland, the Netherlands) and from three populations consisting of three nest-sites in Gemert and Handel (Province of Noord-Brabant, the Netherlands) and Maastricht (Province of Limburg, the Netherlands) (Fig. 1; Table S1). Altogether, in 148 of 149 net-collected individuals we successfully amplified and sequenced 644 bp of the COI gene and 147 were correctly identified as *A. flavipes* (Table S1). The COI sequence of individual 7Caa returned a match with *Stylops melittae*, a Strepsipteran parasite of various *Andrena* species. A total of six unique COI haplotypes were identified in our data set (Fig. S3). A single male (Ned2) net-collected from the Nederrijn, Wageningen, was used for *de novo* genome assembly (Fig. 1; Fig. 2). Nanopore sequencing resulted in 847,773 reads, with a mean read length of 6,192 bp. After assembly, our draft genome consisted of 1600 contigs, with an N50 of 2,442.34 kbp, a total length of 296.6 Mbp and a mean coverage of 16X. BUSCO analysis indicated a highly complete genome (Complete BUSCOs: 99.7 % [Single copy: 99.5 %, Duplicated: 0.2 %], Fragmented: 0.2 %, Missing: 0.1 %, n=1013, E: 13.6 %). Subsequently, the primer pair VgAndrenaF3 and VgAndrenaR15 was designed based on a BLAST search of the *de novo* assembled genome to amplify the putative full-length Vg gene (6072 bp including introns) of *A. flavipes* (Table 1).

In total, the full-length Vg gene was successfully sequenced in 144 out of 148 bees, including for individual 7Caa, for which we could not amplify the *Andrena flavipes* COI. As the Vg sequence of individual 7Caa was identical to that of four other *A. flavipes* individuals in our dataset, individual 7Caa was still identified as *A. flavipes*. Overall, this resulted in 177 Vg sequences of 6072 bp consisting of 111 sequences from males (N=111 haploid males) and 66 from females (N=33 diploid females). After annotation, the full-length of the putative Vg gene was predicted to be 5874 bp, with all amplified 177 Vg sequences having the same coding sequence length, comprising 7 exons and 6 introns (Fig. 3). A total of 78 SNPs were identified in the putative Vg gene of *A. flavipes,* 55 synonymous and 23 non-synonymous, resulting in 85 unique Vg haplotypes (Table S2). Deviations from Hardy-Weinberg equilibrium were detected in several SNPs per population before correction (Table S3A), but none remained significant after Bonferroni correction (Table S3B). Furthermore, linkage disequilibrium (LD) was calculated between each pair of SNP loci in the Vg gene and was generally low (Fig. S4), therefore all SNP loci were retained for further population genetic analyses. Nucleotide diversity (π), Watterson’s theta (θ_W_), number of segregating sites (S), average number of nucleotide differences (K) and Tajima’s D were calculated based on the Vg gene for males and females in each population (Table 2). None of the Tajima’s D values were significantly different from zero, suggesting the observed variation is consistent with neutral evolution, supporting the suitability of the Vg gene for population genetic analyses.

**Figure 3.**
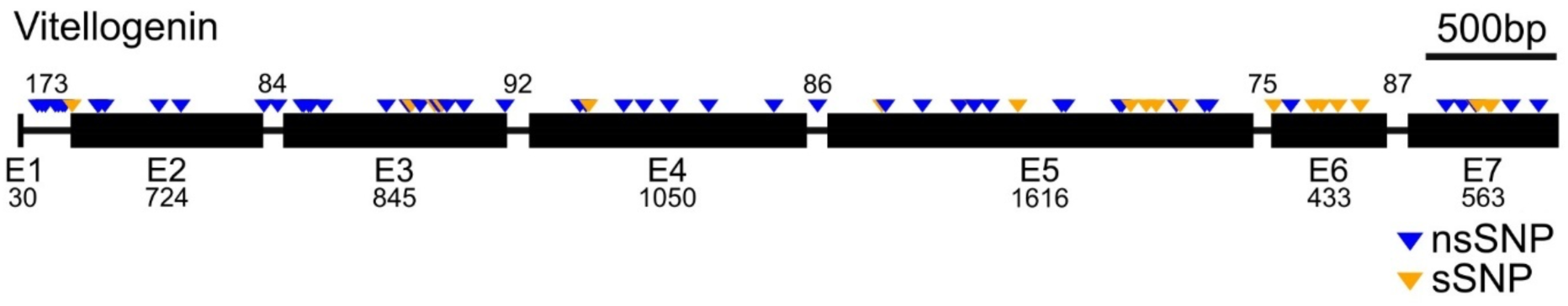
Gene structure diagram with predicted exons and introns of the putative nuclear Vg gene of *A. flavipes* (5874 bp; 78 SNPs). The arrows depict the position of SNPs. In blue are non-synonymous SNPs (nsSNPs) and in yellow are synonymous SNPs (sSNPs). The exons are shown by black boxes and numbered from E1 to E7. The introns are shown by straight black lines. Numbers indicate the size (bp) of the introns (above gene structure diagram) and the exons (below gene structure diagram).

**Table 2.** Genetic diversity parameters for five populations from the Southern Gelderse-Vallei in the Netherlands based on the putative nuclear Vg gene (5874 bp; 78 SNPs) of males and females of *A. flavipes*. *N*=number of Vg sequences; π =nucleotide diversity; *θ_W_ =* Watterson’s theta*; S*=number of segregating sites; *K*=average number of nucleotide differences, *D*= Tajima’s D, 95% CI of D=95% confidence interval of Tajima’s D. Values in brackets indicate the lower and upper bounds [lower, upper] of the 95% confidence interval of Tajima’s D.

| Population | N | S | $\pi$ | $\theta_w$ | K | D | 95% CI of D |
| --- | --- | --- | --- | --- | --- | --- | --- |
| Nederrijn | 85 | 70 | 0.00247 | 0.00238 | 14.487 | 0.124 | [-1.310,1.329] |
| WURCampus | 33 | 54 | 0.00247 | 0.00227 | 14.538 | 0.341 | [-1.332,1.233] |
| Jufferswaard | 30 | 46 | 0.00249 | 0.00198 | 14.634 | 0.966 | [-1.305,1.128] |
| Palmerswaard | 17 | 49 | 0.00243 | 0.00247 | 14.265 | -<br>0.066 | [-1.333,1.286] |
| BlauweKamer | 6 | 37 | 0.00311 | 0.00276 | 18.267 | 0.811 | [-1.214,1.295] |
| All | 171 | 78 | 0.00251 | 0.00232 | 14.767 | 0.255 | [-1.261,1.150] |

As males are haploid and females are diploid, only females could be used to calculate the observed heterozygosity (H_o_), expected heterozygosity (H_e_) and inbreeding coefficient (F_IS_) in our populations. We measured these summary statistics for four *A. flavipes* populations in Southern Gelderse-Vallei, excluding BlauweKamer where only a single female was collected (Table 3). The mean observed heterozygosity per population (H_o_) ranged from 0.179 (Nederrijn) to 0.252 (Palmerswaard) and the mean expected heterozygosity (H_e_) ranged from 0.239 (WURCampus) to 0.282 (Nederrijn), respectively. The mean inbreeding coefficient values (F_IS_) per population ranged from low in the Palmerswaard (-0.051) to very high in the Nederrijn (0.363). The positive F_IS_ values were significantly different from zero in all populations with five or more individuals (Nederrijn, WURCampus and Jufferswaard), suggesting the occurrence of non-random mating in these three populations

**Table 3.** Genetic diversity parameters for four populations from the Southern Gelderse-Vallei in the Netherlands based on the putative Vg gene (5874 bp; 57 SNPs) of females (N=29) of *A. flavipes*. N=number of diploid females collected at population; *H_o_*=observed heterozygosity; *H_e_* =expected heterozygosity; *F_IS_*=inbreeding coefficient; 95% CI of *F_IS_* =95% confidence interval of *F_IS_*. The mean value of *H_o_*, *H_e_* and *F_IS_* are provided per population. Values in brackets indicate the lower and upper bounds [lower, upper] of the 95% confidence interval of *F_IS_*. Significant *F_IS_* values are indicated with an asterisk.

| Population | N | $H_o$ | $H_e$ | $F_{IS}$ | 95% CI of $F_{IS}$ |
| --- | --- | --- | --- | --- | --- |
| Nederrijn | 9 | 0.179 | 0.282 | 0.363* | [0.285, 0.478] |
| WURCampus | 11 | 0.198 | 0.239 | 0.179* | [0.088, 0.281] |
| Jufferswaard | 6 | 0.219 | 0.257 | 0.133* | [0.061, 0.255] |
| Palmerswaard | 3 | 0.251 | 0.252 | -0.051 | [-0.254, 0.253] |

The pairwise *F_ST_* comparisons between each of the five populations in the Southern Gelderse-Vallei were generally low but mostly significant (Table 4). The highest pairwise *F_ST_* value was observed between the Palmerswaard and Jufferswaard (0.055), while the lowest *F_ST_* values were found for most comparisons involving the BlauweKamer (-0.001 to 0.028). All pairwise comparisons involving the BlauweKamer were non-significant, likely due to the limited sample size for this population (N=6). Furthermore, a strong pattern of IBD was found between the five populations tested in the Southern Gelderse-Vallei (Mantel’s r=0.826; P=0.042; Fig. 5), but not when including the outgroup populations. In this scenario, only a moderate and non-significant pattern of IBD was found (Mantel’s r=0.607; P=0.122; Fig. S5).

**Figure 5.**
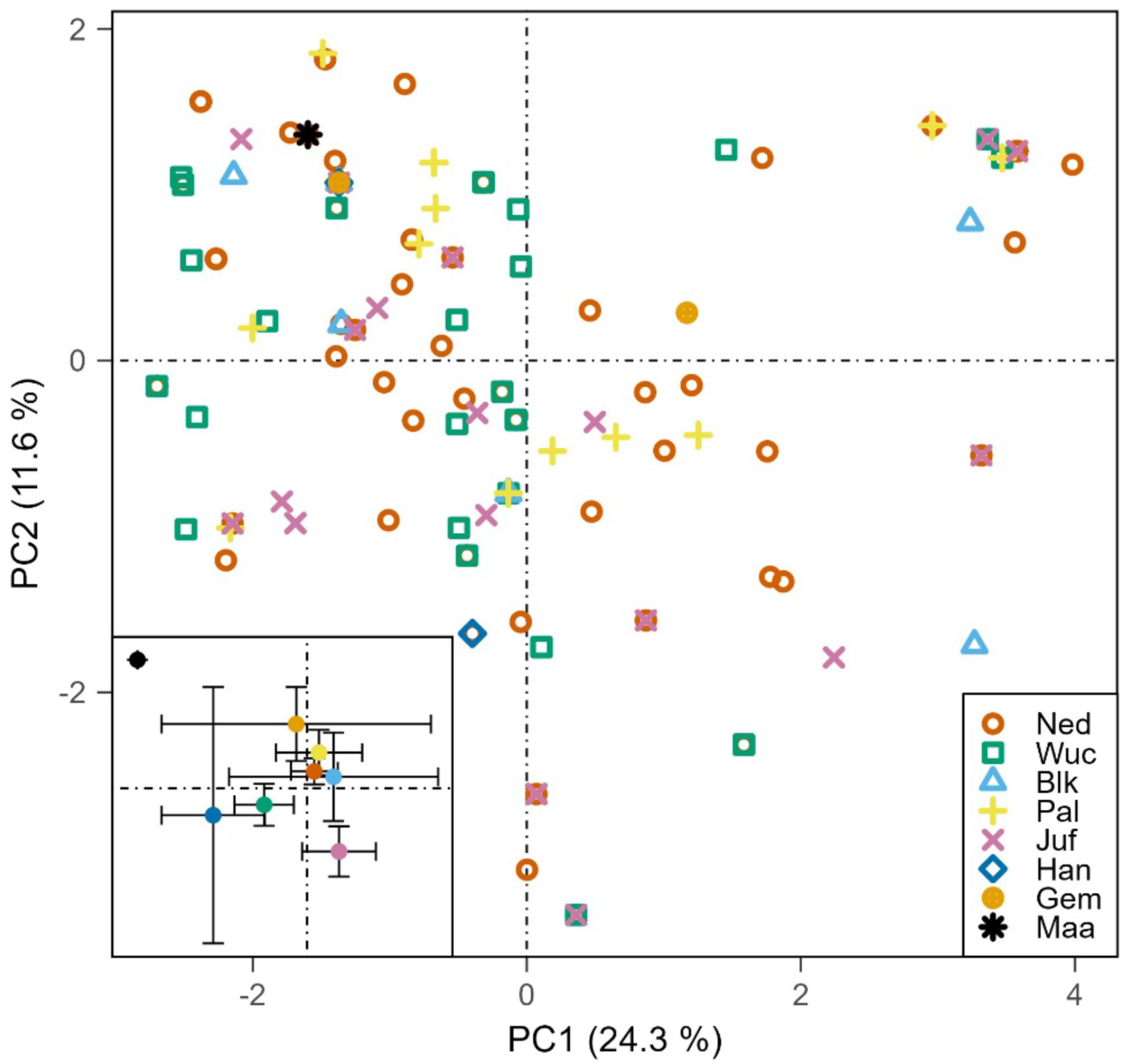
Isolation-by-distance plot of Slatkin Linear’s F_ST_ correlated with geographical (Euclidean) distance (km) in pairwise comparisons between five populations from the Southern Gelderse-Vallei in the Netherlands based on the putative nuclear Vg gene (5874 bp; 78 SNPs) of males and females of *A. flavipes.* Each point represents the average pairwise genetic distance between two populations. Mantel’s r and P-values are given in the plot.

**Table 4.** Pairwise *F_ST_* comparisons between five populations from the Southern Gelderse-Vallei in the Netherlands based on the putative Vg gene (5874 bp; 78 SNPs) of males and females of *A. flavipes*. F_ST_ values are below the diagonal and P-values are above the diagonal. *Ned*=Nederrijn; *Wuc*=WURCampus; *Blk*=BlauweKamer; *Pal*=Palmerswaard; *Juf*=Jufferswaard.

|  | Nederrijn | WURCampus | BlauweKamer | Jufferswaard | Palmerswaard |
| --- | --- | --- | --- | --- | --- |
| Ned | - | 0.027 | 0.404 | 0.008 | 0.000 |
| Wuc | 0.010 | - | 0.421 | 0.000 | 0.010 |
| Blk | 0.003 | 0.002 | - | 0.185 | 0.482 |
| Juf | 0.018 | 0.037 | 0.028 | - | 0.002 |
| Pal | 0.034 | 0.023 | -0.001 | 0.055 | - |

The AMOVA performed with the five populations in the Southern Gelderse-Vallei showed that 2.11 % (*F_ST_* = 0.02114; P<0.001) of the total genetic variation is found between populations and 97.89 % of the total genetic variation within the populations (Table 5). Inclusion of the outgroup populations did not change this pattern, where 2.83 % (*F_ST_* = 0.02833; P<0.001) of the total variation is found between populations, and 97.17 % of the total genetic variation within the populations (Table S4). The Principal Component Analysis (PCA) did not reveal distinct genetic clusters, and the points within each population were widely spread (Fig. 4), confirming that most of the genetic variation is found within and not between the populations of *A. flavipes*. The centroid of each population in the PCA showed that there is a minimal difference between populations, in particular for the outgroup Maastricht population, which is positioned further away from the other populations.

**Figure 4.**
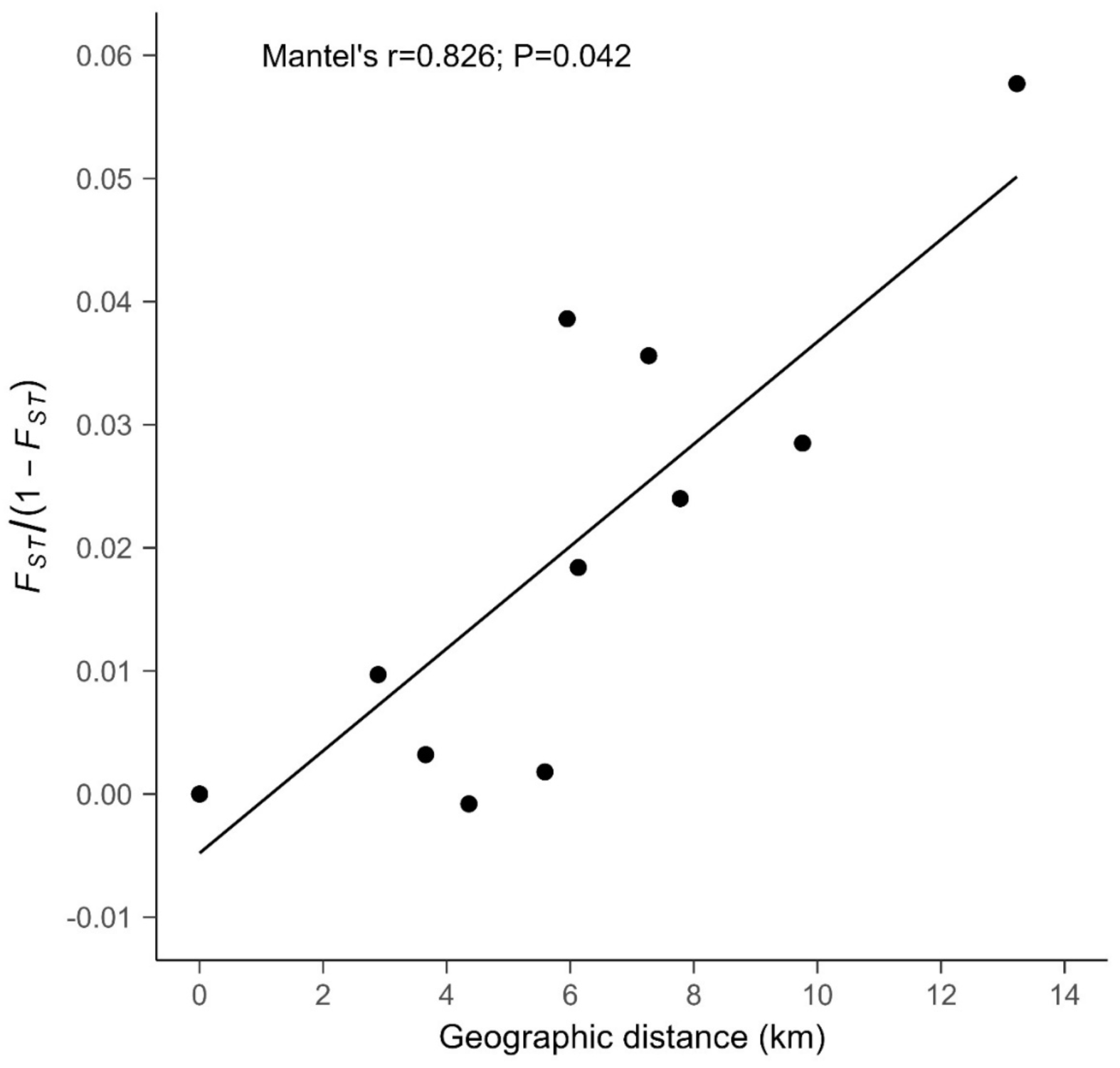
Principal Component Analysis (PCA) of five populations from the Southern Gelderse-Vallei and three outgroup populations in the Netherlands based on the putative nuclear Vg gene (5874 bp; 78 SNPs) from males and females of *A. flavipes*. Each coloured symbol represents a haploid individual, where females are represented as two haploid individuals and males as one haploid individual. The centroids with standard error calculated for each population are given in the lower left panel*. Ned*=Nederrijn; *Wuc*=WURCampus; *Blk*=BlauweKamer; *Pal*=Palmerswaard; *Juf*=Jufferswaard; *Han*=Handel; *Gem*=Gemert en *Maa*=Maastricht.

**Table 5.**
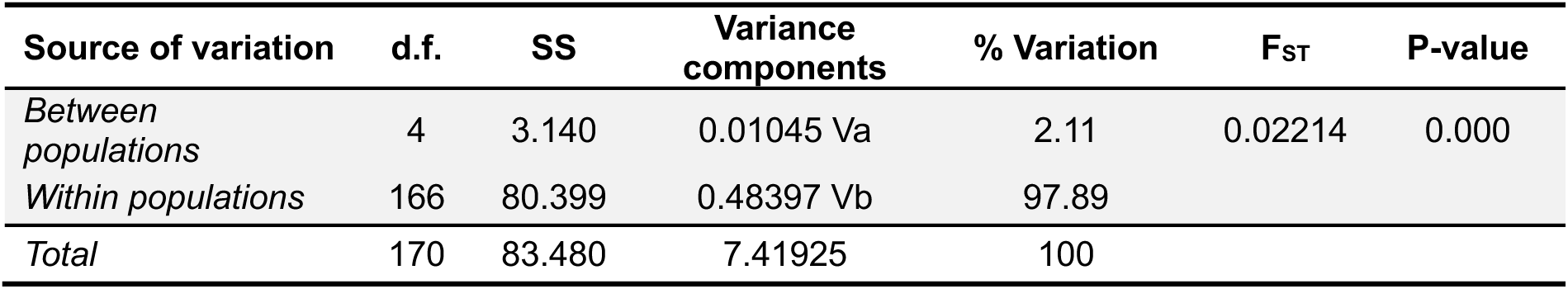
Analysis of Molecular Variance (AMOVA) of five populations from the Southern Gelderse-Vallei in the Netherlands based on the putative Vg gene (5874 bp; 78 SNPs) of males and females of *A. flavipes*. d.f = degrees of freedom; SS=Sum of squares, Va = variance between populations; Vb=variance within populations *F_ST_* = measure of genetic differentiation between populations based on allele frequencies.

## Discussion

Our results reveal low but significant genetic differentiation between *A. flavipes* populations at a local scale (<13 km) in the Netherlands, no distinct genetic clusters, and a pattern of IBD, alongside substantial genetic variation within populations despite heterozygote deficits and positive F_IS_ values in most populations. Together these findings suggest extensive but spatially structured gene flow between nest-aggregations, consistent with non-random mating driven by nest-philopatry. Below we discuss the possible drivers of these patterns, compare them to *A. flavipes* studies at larger spatial scales and to other *Andrena* species, and consider the conservation implications.

### Genetic diversity and inbreeding

In three out of four *A. flavipes* populations studied in the Southern Gelderse-Vallei, a heterozygote deficit and a positive F_IS_ value were detected. Similarly, two population genetic studies of *A. flavipes* in the Iberian Peninsula (de Sousa 2021; de Sousa et al. 2023) and Portugal (Henriques 2021) also reported a heterozygote deficit and positive F_IS_ values. A deficit of heterozygotes and positive F_IS_ within populations may result from population structure (Wahlund effect), non-random mating, or specifically in Hymenoptera their haplodiploid genetic system. The heterozygote deficit and a positive F_IS_ observed for *A. flavipes* is unlikely due to population structure, as the PCA, AMOVA and F_ST_ analysis revealed low genetic differentiation and no clustering between populations. Consistently, in the Iberian Peninsula, populations were first inferred from the data rather than assigned a priori, after which F_IS_ was measured within each group, showing that their positive F_IS_ values were not due to a Wahlund effect (de Sousa et al. 2023).

The heterozygote deficit and high positive F_IS_ values in *A. flavipes* might stem from its haplodiploid genetic system, in which males are haploid and females are diploid. Therefore, haplodiploids have fewer gene copies per generation than diplodiploids, lowering N_e_. This increases the effects of genetic drift and inbreeding, which can lead to a heterozygote deficit and a positive F_IS_ (Packer & Owen 2001). Additionally, due to bees’ complementary sex determination system, inbreeding can lead to diploid male production, further reducing N_e_ and increasing the effects of genetic drift and inbreeding (Zayed & Packer 2005). In many wild bee species, a heterozygote deficit and positive F_IS_ has been reported (Nogueira-Neto 2002; Danforth et al. 2003; Zayed et al. 2005; Darvill et al. 2006; Stow et al. 2007; Zayed & Packer 2007; Kent et al. 2018; Harpur & Rehan 2021; Samad-zada et al. 2023), including other *Andrena* species, namely *Andrena scotica, A. fuscipes, A. vaga, A. agilissima, A. haemorrhoa* and *A. nigroaenea* (Paxton et al. 1996; Exeler et al. 2008, 2010; Černá et al. 2013; de Sousa et al. 2023; Tomowski et al. 2025).

Although haplodiploidy cannot be excluded as a contributor to the observed heterozygote deficit and positive F_IS_, non-random mating due to communal nesting behaviour is likely the more important driver of inbreeding in *A. flavipes*. This also appears true for the other *Andrena* species, of which only *A. fuscipes* does not nest communally and indeed showed lower inbreeding values compared to communal nesting *Andrena* species (Exeler et al. 2010). *A. flavipes* females often nest in large aggregations, and they tend to only mate and nest within their natal nest-aggregation. Furthermore, males have been found to prefer to mate with females within their own natal nest-aggregation (Butler 1965). This nest-site philopatric tendency has been hypothesized to promote non-random mating in bees (Paxton 2005; López-Uribe et al. 2015), which is reflected in the positive F_IS_ values observed in *A. flavipes*. Moreover, within *A. scotica* populations, high inbreeding has been directly linked to intranidal mating, where mating occurs inside the natal nests prior to females’ emergence in spring (Paxton & Tengö 1996). Microsatellite analysis revealed very low diploid male production in *A. scotica* (Paxton et al. 2000), suggesting that the haplodiploid system alone is not sufficient to explain the heterozygote deficit and high inbreeding in this species, and that most likely its nesting behaviour plays a role. This low diploid male production may partly reflect multi-locus CSD in *Andrena* rather than the single-locus CSD of eusocial bees (Leung & van der Meulen, 2022), which substantially lowers the probability of producing diploid males through inbreeding. Consistent with a role for nesting behaviour rather than haplodiploidy alone, high positive F_IS_ values were also found for *A. vaga* (Exeler et al. 2008; Černá et al. 2013), possibly related to its philopatric nesting habits, in which males tend to wait at nest-sites to mate with freshly emerged females (Vleugel 1947; Hallmen 1991) and females are philopatric (Westrich 1989b). Based on the biology of the *Andrena* genus, the observed heterozygote deficit and positive F_IS_ in *A. flavipes* are most likely due to its reported nest-philopatry and potential high frequency of within-nest mating.

Despite evidence of high inbreeding, *A. flavipes* in the Southern Gelderse-Vallei showed substantial genetic variation within its populations. Likewise*, A. flavipes* populations in Austria and the Loire Valley in France showed high genetic variation (Villalta et al. 2021; Rupprecht 2022). Interestingly, also in populations of *A. fuscipes*, *A. vaga*, *A. nigroaenea* and *A. haemorrhoa*, inbreeding did not lead to reduced genetic diversity (Exeler et al. 2008, 2010; Tomowski et al. 2025). High genetic diversity despite inbreeding in *Andrena* species might be explained by a relatively large effective population size (N_e_), resulting from reproduction by all females (Romiguier et al. 2014). Unlike eusocial Hymenoptera where only a small number of queens contribute to reproduction, all females in *Andrena* species breed, which partially offsets the N_e_-reducing effect of haplodiploidy. Furthermore, a female-biased sex ratio in many *Andrena* species (Paxton & Tengö 1996; Paxton et al. 1999; Bischoff 2003; Maeta et al. 2004; Lund Norbakk 2017; Sirohi et al. 2025) increases N_e_ relative to diplodiploids with the same census size (Packer & Owen 2001). Moreover, as males are haploid, they express deleterious alleles directly, allowing selection to effectively purge them from the population (Brückner 1978; Werren 1993; Antolin 1999), lowering the genetic load and the potential impact of inbreeding in haplodiploids.

### Population genetic structure

The weak local population structure in *A. flavipes* is likely due to its high abundance and widespread distribution in Europe (Alfken 1913; Hallmen 1993; Peeters et al. 2012), which increases the likelihood of genetic exchange between nest-aggregations, despite its observed philopatric behaviour. Furthermore, while nest-aggregations of *A. flavipes* can persist for several decades (Perkins 1919; Boyes et al. 2024), most are likely short-lived, with frequent (re)colonization, population fluctuations and local extinction events, as demonstrated in Zeeland, the Netherlands and Bedfordshire, UK (Chambers 1968; Calle & Dobbelaar 2007), resulting in the observed weak population structure. These transient colonization and extinction patterns, together with the occasional long-term persistence of some populations, also observed in *Andrena humilis* and *A. vaga* (Perkins 1919; Ulrich 1956; Bischoff 2003; Fellendorf et al. 2004; Franzén & Nilsson 2013), support metapopulation dynamics across *Andrena* species, with persistent populations acting as sources for temporary sink populations, as demonstrated for *A. humilis* (Franzén & Nilsson 2013).

Studies investigating the population genetic structure of *A. flavipes* have revealed that strong population structuring occurs mainly at national and continental scales, while local scales (<11 km) show weak or no differentiation. This was demonstrated by Ortego et al. (2024), who in a European study found only *A. flavipes* populations in Denmark to show weak but significant differentiation (F_ST_=0.03), while this was lacking in populations from the Netherlands, Switzerland and Spain. Regional to national-scale analyses in Austria (25–400 km) and Portugal (10–250 km) similarly found no population structure (Henriques 2021; Rupprecht 2022). However, expanding the scale to include Spain (<800 km) and continental Europe (400–2,500 km) uncovered distinct populations: two highly differentiated clusters in the Pyrenees and Iberia (de Sousa 2021; de Sousa et al. 2023), and three populations in Denmark, Spain, and Central Europe (Netherlands/Switzerland), with Spain exhibiting the highest genetic differentiation, while the Denmark cluster only became apparent after excluding Spain (Ortego et al. 2024). Moreover, IBD was detected at this scale (Ortego et al. 2024). This suggests that the Pyrenees population identified by de Sousa et al. (2023) may belong to the Central European cluster. The strong differentiation of the Iberian Peninsula is consistent with historical isolation in glacial refugia, whereas the emergence of a northern cluster (Denmark) is more likely driven by recent range expansion (Cederberg & Nilsson 2000; Edwards 2002; Rasmussen et al. 2016), creating a founder effect (Ortego et al. 2024).

Other *Andrena* species show comparable patterns, such as *A. fuscipes* showing weak genetic differentiation and no IBD at local to regional scales in Germany (4 - 150 km; Exeler et al. 2010), while *A. haemorrhoa* and *A. nigroaenea* showed weak differentiation and no correspondence between genetic clustering and spatial clustering at local scales (70 m - 15.6 km; Tomowski et al. 2025). *A. vaga* showed no genetic differentiation nor IBD at regional scales in the Upper-Rhine Valley (400 m - 75 km; Mohra et al. 2004), but weak genetic differentiation and IBD at the national scale in the Lower Rhine Valley (750 m - 330 km; Exeler et al. 2008), and distinct Western and Eastern populations at regional and national scales in the Czech Republic (25 - 400 km; Černá et al. 2013). Similarly, *A. agilissima* showed three highly differentiated populations in the Iberian Peninsula (<800 km; de Sousa 2021; de Sousa et al. 2023). Notably, most of these studies relied on microsatellite markers, and it is possible that marker choice and number influence the detection of fine-scale structure; future genome-wide studies would be needed to determine whether IBD at local scales is consistently detectable across philopatric *Andrena* species. Overall, these findings suggest that *Andrena* species may generally lack strong genetic structure at local to regional scales in Europe, but exhibit structuring at larger, more national to continental scales, emphasizing the influence of spatial scale and biogeography on population genetic patterns.

### Conservation implications

Our results show a weak but significant population structure and substantial genetic variation within populations of *A. flavipes*, suggesting that gene flow between populations is currently maintained, likely facilitated by *A. flavipes’* high abundance (Alfken 1913; Hallmen 1993; Peeters et al. 2012; Kleijn et al. 2015). Our results also show high F_IS_ values and a pattern of IBD, indicating non-random mating and spatially structured gene flow. While gene flow appears currently sufficient to prevent strong differentiation, increasing spatial isolation could lead to stronger differentiation over time. A reduction in connectivity between nest-aggregations may be particularly important given the metapopulation dynamics suggested for *A. flavipes*, as it could limit the extent to which larger populations contribute to the persistence of smaller populations that are more prone to local extinction, increasing the impact of local population loss on overall connectivity within the metapopulation.

Maintaining connectivity between nest-aggregations should therefore be prioritized in the management of the species. This is of particular concern because the species has been shown to be less abundant in more urbanized landscapes (Fischer et al. 2016) and to decline with increasing agricultural land-use intensification (Kleijn et al. 2015; Ockermüller et al. 2023). In agricultural landscapes, semi-natural elements such as grassy field margins and herbaceous strips function as source habitats for bees (Krewenka et al. 2011), supporting their movement from these habitats into adjacent agricultural fields. Specifically, for *A. flavipes*, higher gene flow from semi-natural habitats into agricultural landscapes has been found (Ortego et al. 2024), suggesting that semi-natural habitats act as reservoirs of genetic diversity for *A. flavipes*. Conservation efforts should therefore focus on the preservation and restoration of semi-natural habitats across agricultural and urbanized landscapes to maintain landscape connectivity and thereby facilitate gene flow between nest-aggregations.

In addition to connectivity, the habitat quality around and within the nest-aggregations is also likely to be important in the long-term persistence of nest-aggregations. *A. flavipes* tends to be larger with increasing distance from semi-natural landscapes (Warzecha et al. 2016). Since dispersal range in bees scales with body size (Greenleaf et al. 2007; López-Uribe et al. 2019), this suggests that individuals in more intensively managed agricultural landscapes may need to travel further to meet their floral resource requirements, implying that local floral availability is reduced in these settings. Floral resource availability may be especially important for local population persistence, as higher pollen abundance in other andrenid species *A. humilis*, *Andrena hattorfiana* and *Andrena clarkella* has been linked to increased survival of local populations (Calle & Dobbelaar 2002; Franzén & Nilsson 2010, 2013) and to the formation of large nest-aggregations in *A. vaga* (Westrich 1989a). Suitable nesting habitat is also crucial: dense vegetation cover contributes to the extinction of local A. flavipes populations (Chambers 1968; Calle & Dobbelaar 2007). Extensively managed meadows with infrequent mowing, no fertilization, lower vegetation height and small patches of bare ground support higher A. flavipes abundances than intensively managed meadows (Albrecht et al. 2023). Overall, effective conservation of *A. flavipes* in human-altered landscapes will require maintaining both landscape connectivity, facilitated by the restoration and preservation of semi-natural habitats, as well as the availability of sufficient floral resources and suitable nesting places at a local scale.

## Acknowledgements

We thank Utrechts Landschap for access permission to collect on their sites. We are grateful to Patrick G.H. Verbaarschot and Francisca C. Reyes Marquez for assistance in the lab, especially with the library preparation and nanopore sequencing. We would like to thank Hendrik-Jan W.C. Megens who assisted with the initial data processing of the Vg reads and visualization of our data. Furthermore, we are grateful to Reindert Nijland, Hylke Kortenbosch, Mees Horssen and Erik van der Spek for the help in collecting *A. flavipes* in the field.

## Statements & Declarations

### Author contributions

Conceptualization: Bart A. Pannebakker; Lotte F. Wösten; Joost van den Heuvel; Frank F.M. Becker; Methodology: Frank F.M. Becker, Lotte F. Wösten, Joost van den Heuvel; Bart A. Pannebakker; Formal analysis and investigation: Lotte F. Wösten; Writing - original draft preparation: Lotte F. Wösten; Writing - review and editing: Bart A. Pannebakker, Joost van den Heuvel, Frank F.M. Becker; Visualization: Lotte F. Wösten; Funding acquisition: Bart A. Pannebakker; Resources: Bart A. Pannebakker, Frank F.M. Becker; Supervision: Bart A. Pannebakker.

### Data Availability

COI and vitellogenin amplicons, and raw sequencing reads were deposited on NCBI in Bioproject PRJNA1468685.

