## Supplementary Figures for "Local population genetic structure of *Andrena flavipes* (Hymenoptera, Andrenidae), a nest-philopatric communal mining bee, reveals isolation-by-distance and high inbreeding"

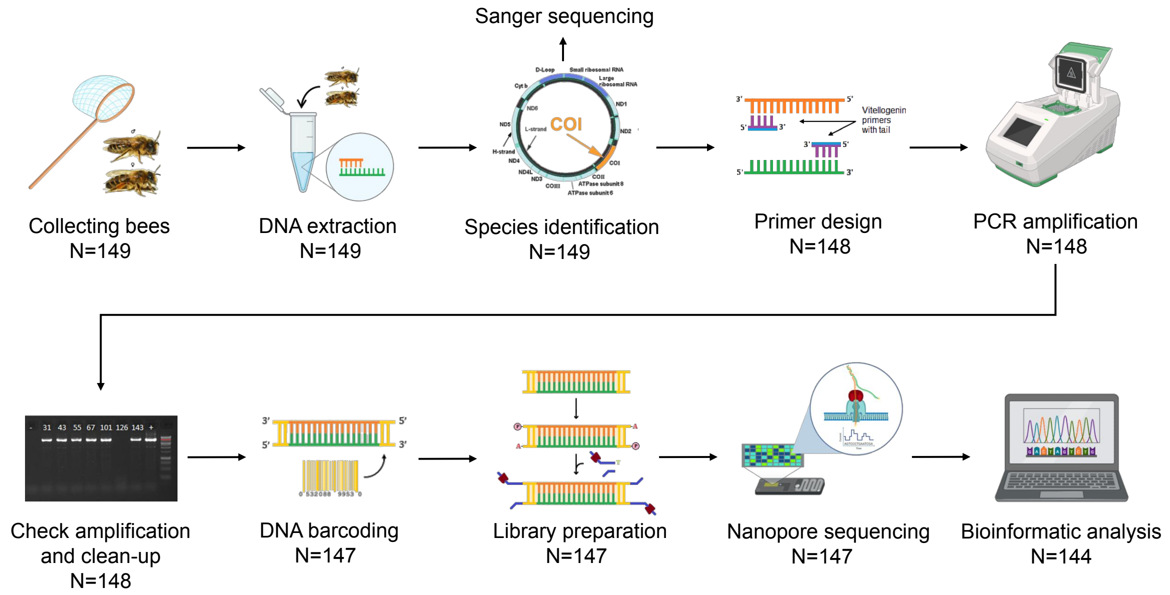


Figure S1. **Schematic overview of methods.** N indicates how many bee samples remain after performing each step. Sequences indicates the number of Vg sequences that were in total obtained.

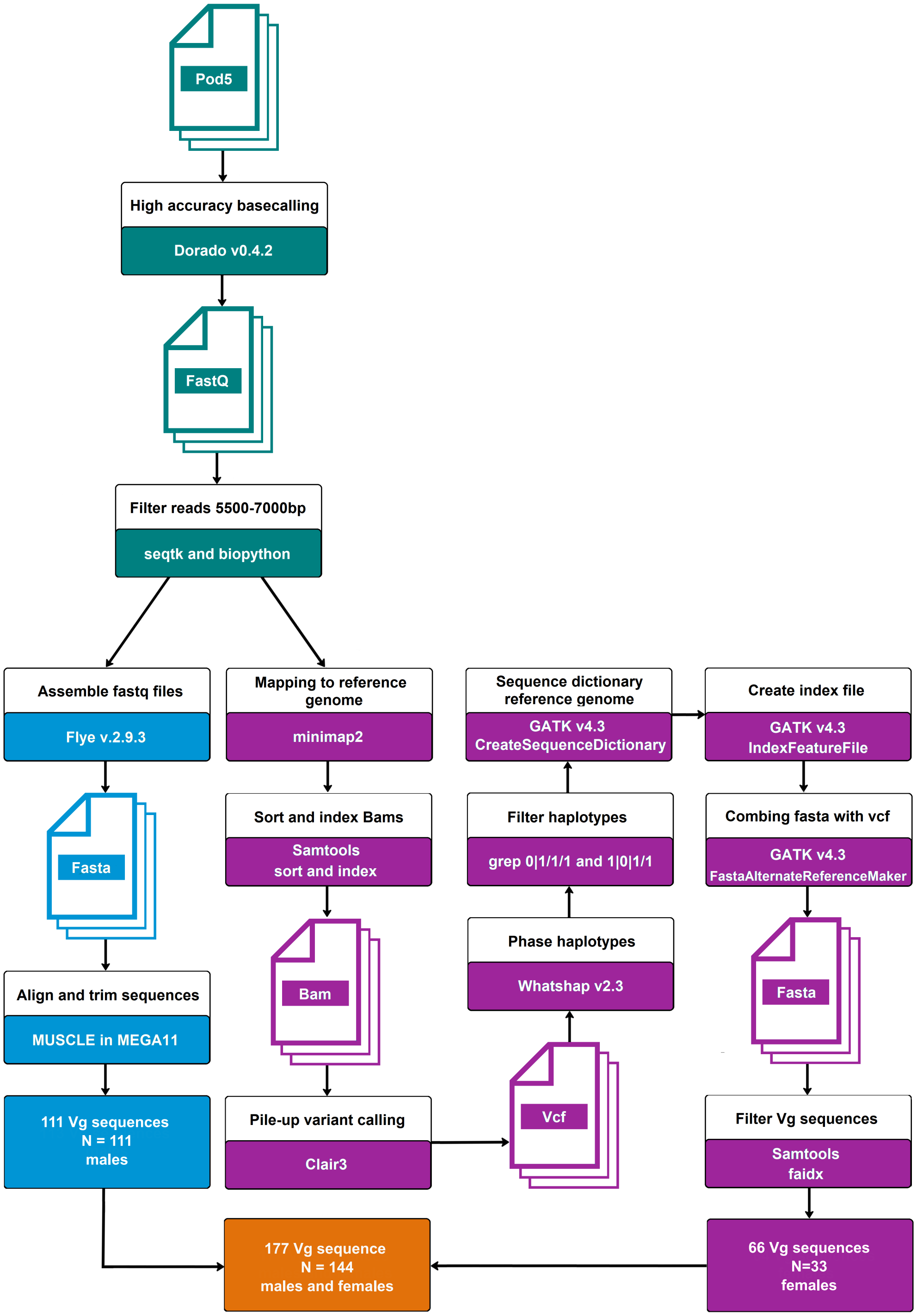


Figure S2**. Schematic overview of bioinformatic pipeline**. In green the pipeline for both males and females, in blue the pipeline for only males and in purple the pipeline for only females. In orange are the total number of Vg sequences combined.

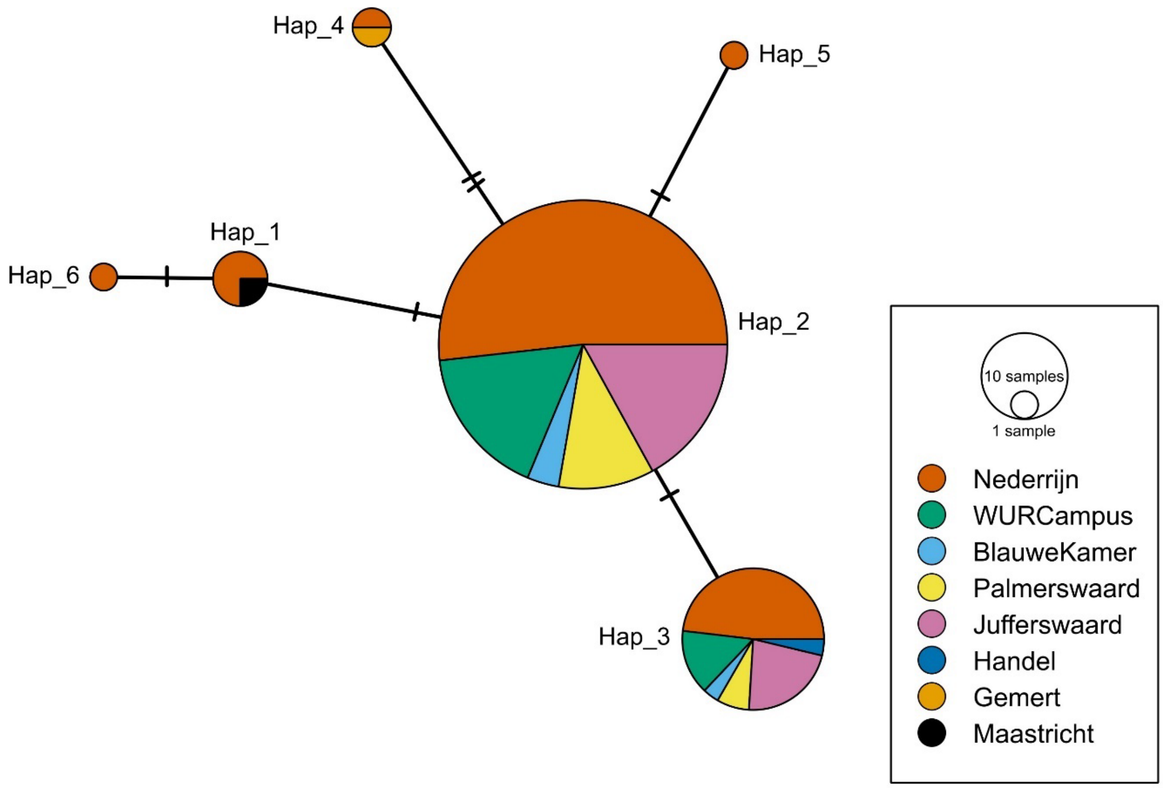


Figure S3. **Median-joining haplotype network of five populations from the Southern Gelderse-Vallei and three outgroup populations in the Netherlands based on part of the COI gene (644 bp) of males and females of *A. flavipes*.** Each coloured circle represents a unique haplotype, and the size of each circle is proportional to the frequency of the haplotype it represents. A dash corresponds to a single mutation between the connected haplotypes. A colour code is given for each population in the legend on the right.

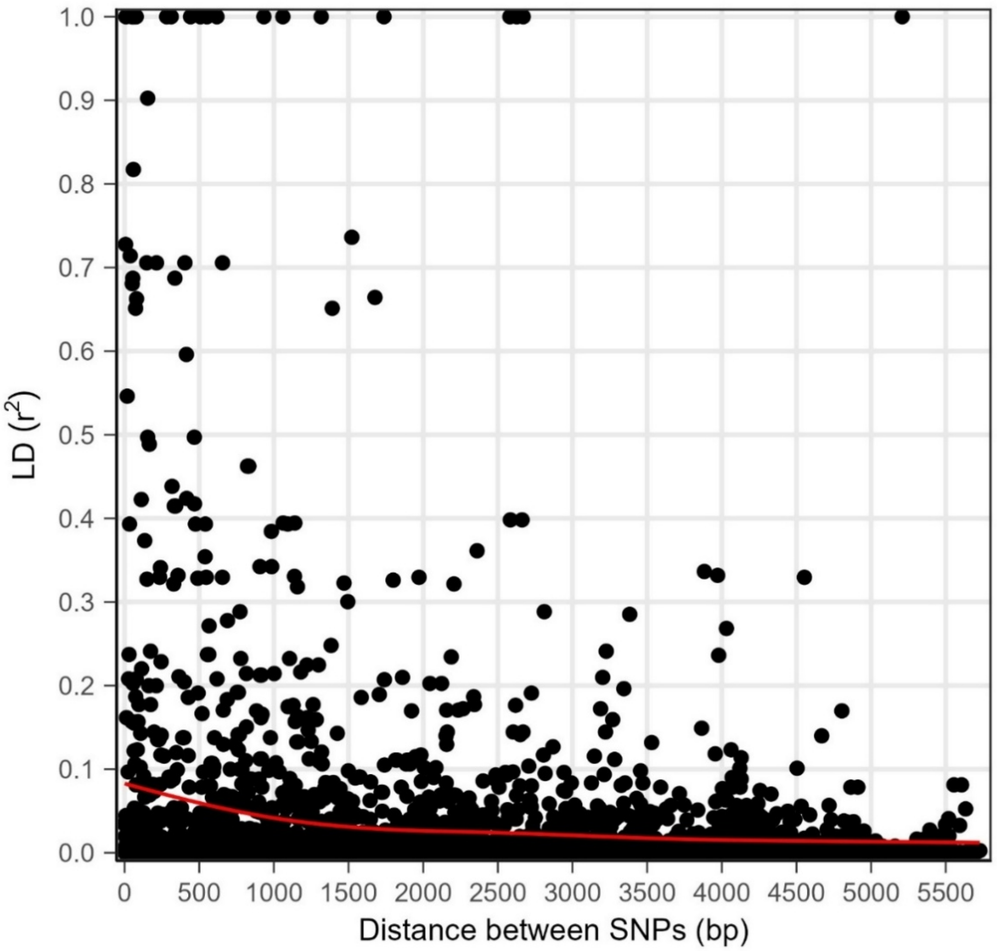


Figure S4. **Linkage disequilibrium decay plot of LD ($r^2$) as a function of genetic distance between SNPs (bp) of the Vg gene of *A. flavipes*.** Grey dots indicate the observed pairwise LD (3003 test total) and the red line shows the average of LD ($r^2$) as windows of 500 bp.

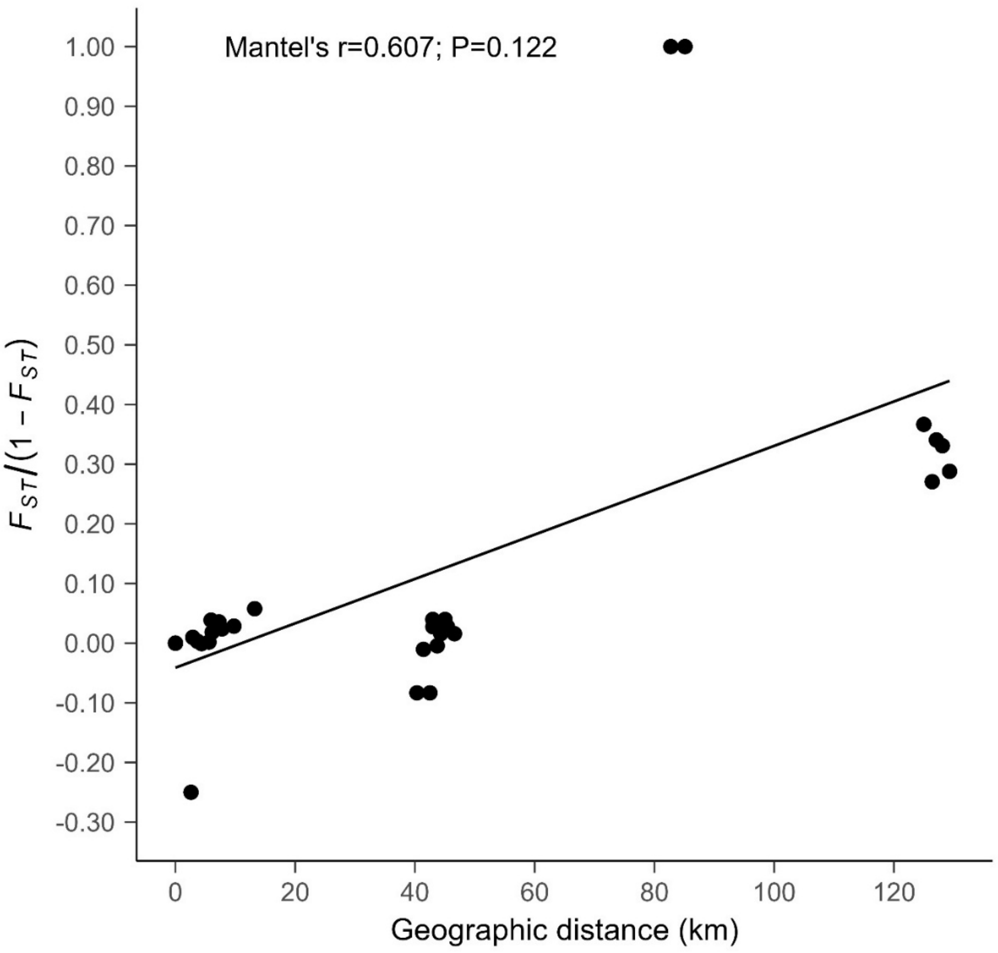


Figure S5. **Isolation-by-distance plot of Slatkin Linear's $F_{ST}$ correlated with geographical (Euclidian) distance (km) in pairwise comparisons between five populations from the Southern Gelderse-Vallei and three outgroup populations in the Netherlands based on the putative nuclear Vg gene (5874 bp; 78 SNPs) of males and females of *A. flavipes.*** Each point represents the average pairwise genetic distance between two populations. Mantel's r and P-values are given in the plot.
